# Gigaxonin is required for intermediate filament transport

**DOI:** 10.1101/2022.08.19.504572

**Authors:** Bhuvanasundar Renganathan, James P Zewe, Yuan Cheng, Mark Kittisopikul, Puneet Opal, Karen M Ridge, Vladimir I. Gelfand

**Affiliations:** Department of Cell and Developmental Biology, Feinberg School of Medicine, Northwestern University, Chicago, IL, United States; Ken and Ruth Davee Department of Neurology, Feinberg School of Medicine, Northwestern University, Chicago, IL, United States; Division of Pulmonary and Critical Care Medicine, Department of Medicine, Northwestern University Feinberg School of Medicine, Chicago, IL, United States

**Keywords:** GAN, intermediate filaments, kinesin-1, dynamics, vimentin, neurofilaments

## Abstract

Gigaxonin is an adaptor protein for E3 ubiquitin ligase substrates. It is necessary for ubiquitination and degradation of intermediate filament (IF) proteins. Giant axonal neuropathy is a pathological condition caused by mutations in the *GAN* gene that encodes gigaxonin. This condition is characterized by abnormal accumulation of IFs in both neuronal and non-neuronal cells; however, it is unclear what causes IF aggregation. In this work, we studied the dynamics of IFs using their subunits tagged with a photoconvertible protein mEOS 3.2. We have demonstrated that the loss of gigaxonin dramatically inhibited transport of IFs along microtubules by the microtubule motor kinesin-1. This inhibition was specific for IFs, as other kinesin-1 cargoes, with the exception of mitochondria, were transported normally. Another effect of gigaxonin loss was a more than 20-fold increase in the amount of soluble vimentin oligomers in the cytoplasm of gigaxonin knock-out cells. We speculate that these oligomers saturate a yet unidentified adapter that is required for kinesin-1 binding to IFs, which might inhibit IF transport along microtubules causing their abnormal accumulation.

## Introduction

Gigaxonin is a ubiquitously expressed protein encoded by the *GAN* gene located on human chromosome 16q23.2 (1, 2). The lack or loss of function of gigaxonin causes giant axonal neuropathy (GAN), an autosomal recessive disorder. The *GAN* gene has more than 50 distinct loss of function mutations that cause the disorder (3–6). The loss of both central and peripheral neurons leads to gradual muscular atrophy in this pathological condition. Patients often begin displaying symptoms between the ages of 3 and 5 years old and die between the ages of 20 and 30 (7).

Gigaxonin is a member of the BTB (Bric-a-brac, Tram track and, Broad) /Kelch domain superfamily (8). E3 ligase cullin3 (Cul3), a member of the 26S proteasomal complex, binds to the BTB domain of gigaxonin, whereas intermediate filaments (IFs) interact with the Kelch domain (9). Thus, gigaxonin functions as a Cul3 ligase adaptor, ubiquitinating IFs and regulating IF turnover via proteasomal degradation (10). *GAN* gene loss causes an aberrant accumulation of neurofilaments (NFs) in neuronal axons, resulting in a giant axonal phenotype (11). In addition to NFs, all other classes of IFs are disorganized and aggregated in non-neuronal cells of GAN patients (12).

IFs are cytoskeletal structures required to stabilize cells against both mechanical and non-mechanical stress (13, 14). In the cytoplasm, IFs scaffold membrane organelles and connect with desmosomes and focal adhesions at the plasma membrane to build a network that holds the nucleus in place. Six types of IFs have been characterized based on their amino acid sequence. The structural organization of IF proteins is homologous, with a central rod domain and varying N- and C-termini (15). NFs are type IV IFs that form heteropolymers comprising three NF proteins (NFL, NFM, and NFH) as well as related IF proteins alpha-internexin and peripherin. NFs stabilize neuronal morphology and participate in axonal development and other important neuronal processes (17). Various neurodegenerative pathologies have been linked to disorganization and loss of functional NFs (14, 17, 18), which result in reduced axonal development, neuronal caliber loss, and impaired transport of organelles and cargos (19, 20). Mutations in genes encoding NF proteins or their metabolism are widely accepted as the cause of disorganized NFs in neurodegenerative disorders (16). GAN falls into the latter group since mutations in the *GAN* gene impede the degradation and organization of all IFs, including NFs. However, a molecular mechanism of IF aggregation in affected cells is unknown.

Normal distribution of IFs in a cell depends on cytoplasmic microtubules (18–20). We and others have shown that IFs are transported along microtubules by the major plus-end directed microtubule motor kinesin-1 (19, 21). In a mouse model, kinesin-1 mutations promote inefficient anterograde transport of NF in neurons, as well as disordered accumulation of NFs in neuronal cell bodies (22). Due to the absence of anterograde transport, IFs are disorganized and aggregate near the nucleus in kinesin-1 knockout cells (23). The similarities in IF aggregation and disorganization patterns between kinesin-1 knockout and GAN suggests that IF aggregation in the cells of GAN patients could be triggered by impaired IF transport along microtubules. To address this possibility, we examined IF transport in *GAN* KO cells by applying fiduciary markers on the IF using a photoconversion approach. We observed that the loss of gigaxonin had a significant negative impact on the transport of IFs, while the motor transporting IFs was fully active and could move other cargoes. We also noticed a substantial increase in the soluble form of vimentin in *GAN* KO cells.

## Results

### *GAN* gene loss inhibits vimentin IF transport

IFs form a dense network extending from the nucleus towards the cell periphery and act as a mechanical stabilizer for the cell. IFs undergo constant rearrangement and transport along microtubules to facilitate the ongoing requirements of cellular activity (24). We hypothesized that inhibition of normal IF dynamics causes IF disorganization in GAN. To test this hypothesis, we studied the transport of the vimentin IFs (VIF) in *GAN* knockout (*GAN* KO) cells using a photoconversion strategy. We used RPE cells stably expressing vimentin fused with a photoconvertible protein mEOS3.2 (mEOS-vim). mEOS-vim copolymerizes with endogenous vimentin and forms normal VIF networks in RPE cells (Fig 1A). Using CRISPR, we knocked out the *GAN* gene in RPE cells stably expressing mEOS-vim. In *GAN* KO cells normal VIF organization was disrupted and the majority of VIFs formed aggregates in the juxtanuclear region, but some labeled filaments were observed at the cell periphery (Fig 1B). This VIF disorganization is a typical phenotype of GAN KO cells (25, 26).

**Figure 1:**
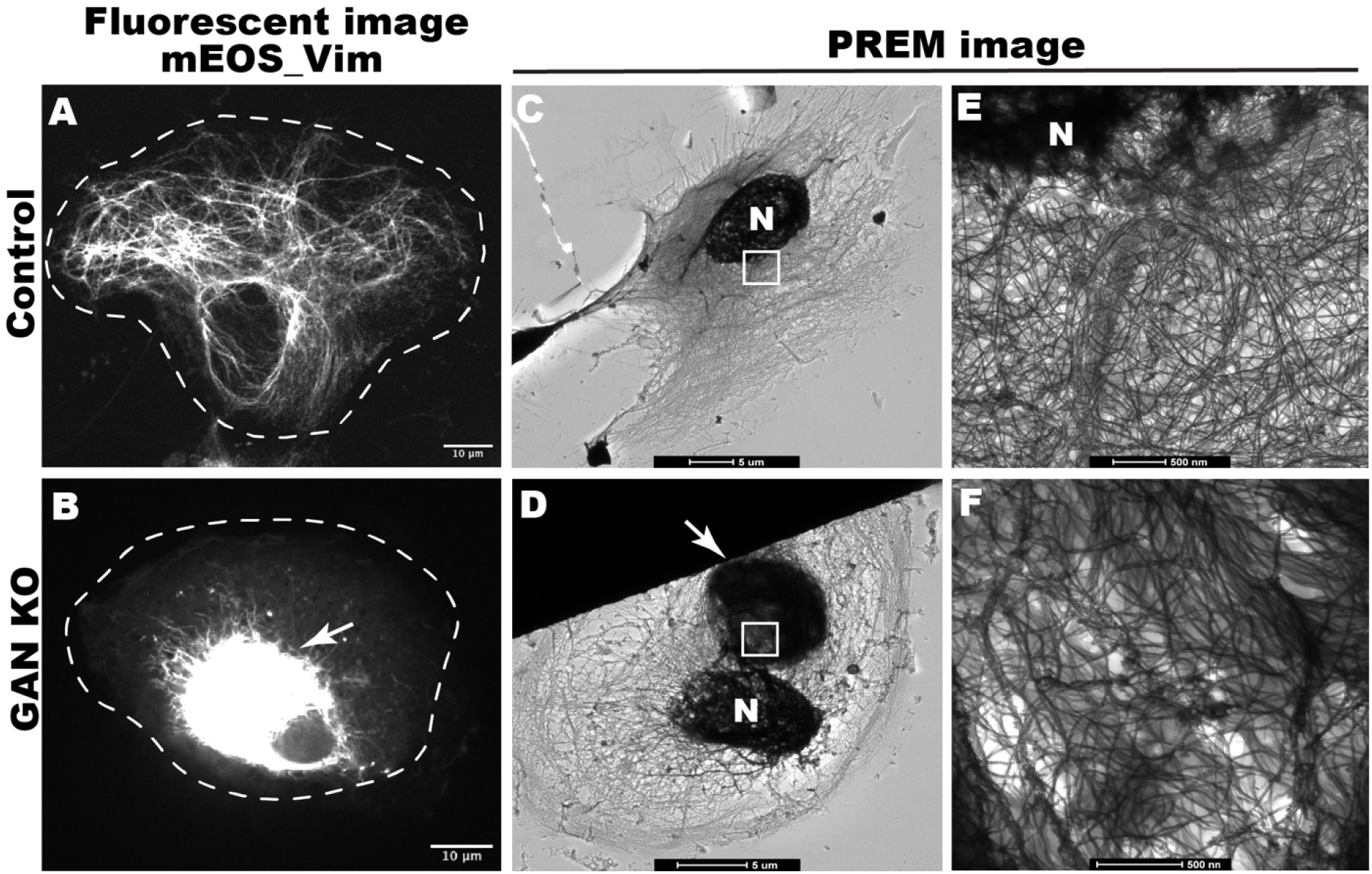
*GAN* KO affects organization of IFs. A) shows the normal distribution of mEOS_vim in control RPE cell, B) shows the mEOS_vim aggregation in *GAN* KO RPE cells. Scale bar 10 micrometers. Platinum replica electron microscope (PREM) images of control cell (C & E) and *GAN* KO cell (D & F). IF meshwork in the entire cell can be appreciated in lower magnification (C) and single filaments can be observed in higher magnification near nucleus (E). In *GAN* KO cells, juxtanuclear IF aggregates can be observed at lower magnification (D), single filaments can be observed in the IF aggregate at higher magnification of aggregated region (F). Nucleus is indicated by N, arrow points to IF aggregates and white box in C & D represent the magnified region for E & F respectively. Scale bar 5 μm for C & D; 500 nm for E & F PREM images.

Platinum replica electron microscopy (PREM) was used for high-resolution analysis of VIF organization and distribution in *GAN* KO cells. In control cells, we observed a fine meshwork of VIFs (Fig 1C). In *GAN* KO cells, we observed dense VIF aggregates in the juxtanuclear position (Fig 1D). At higher magnification, single filaments could be resolved in all regions of the control cell (Fig 1E). In *GAN* KO cell, VIF formed a dense interwoven network and bundles, yet single filaments could be observed within the aggregates (Fig 1F). The presence of single filaments in *GAN* KO aggregates motivated us to examine VIF transport these cells.

We have demonstrated previously that cytoskeletal dynamics can be studied efficiently by fluorescence microscopy of live cells using photoconversion of mEOS fluorescent protein fused with a cytoskeletal protein of interest (20). Emission of mEOS can be photoconverted from the green (488 nm) to the red (561 nm) channel by illumination with a 405 nm laser. Using a pinhole aperture, a small region of mEOS-vim-labeled filaments in a live cell was photoconverted and cells were imaged in the red channel. This technique showed that control cells display robust transport of VIFs away from the photoconverted zone. In control cells, several filaments were typically transported during the total imaging time (three minutes), demonstrating active VIF transport under normal conditions (Fig 2C C’ and Video 1). However, mEOS-vim in *GAN* KO cells behaved differently. All photoconverted filaments remained confined within the initial photoconverted zone throughout the duration of the experiment. These results showed that VIF transport was severely affected by the loss of *GAN* gene (Fig 2F, F’ and Video 2). It is possible that the lack of VIF transport could be a secondary effect of IF aggregation. To test this idea, we photoconverted a region of the cytoplasm in *GAN* KO cells where individual filaments could still be found. We found that even unaggregated VIFs in *GAN* KO cells were practically immotile (Video 3). We have quantified the total length of segments of VIF filaments transported outside of the photoconverted zone in control and *GAN* KO cells. This quantification showed dramatic inhibition of VIF transport in *GAN* KO cells (Fig 2G).

**Figure 2:**
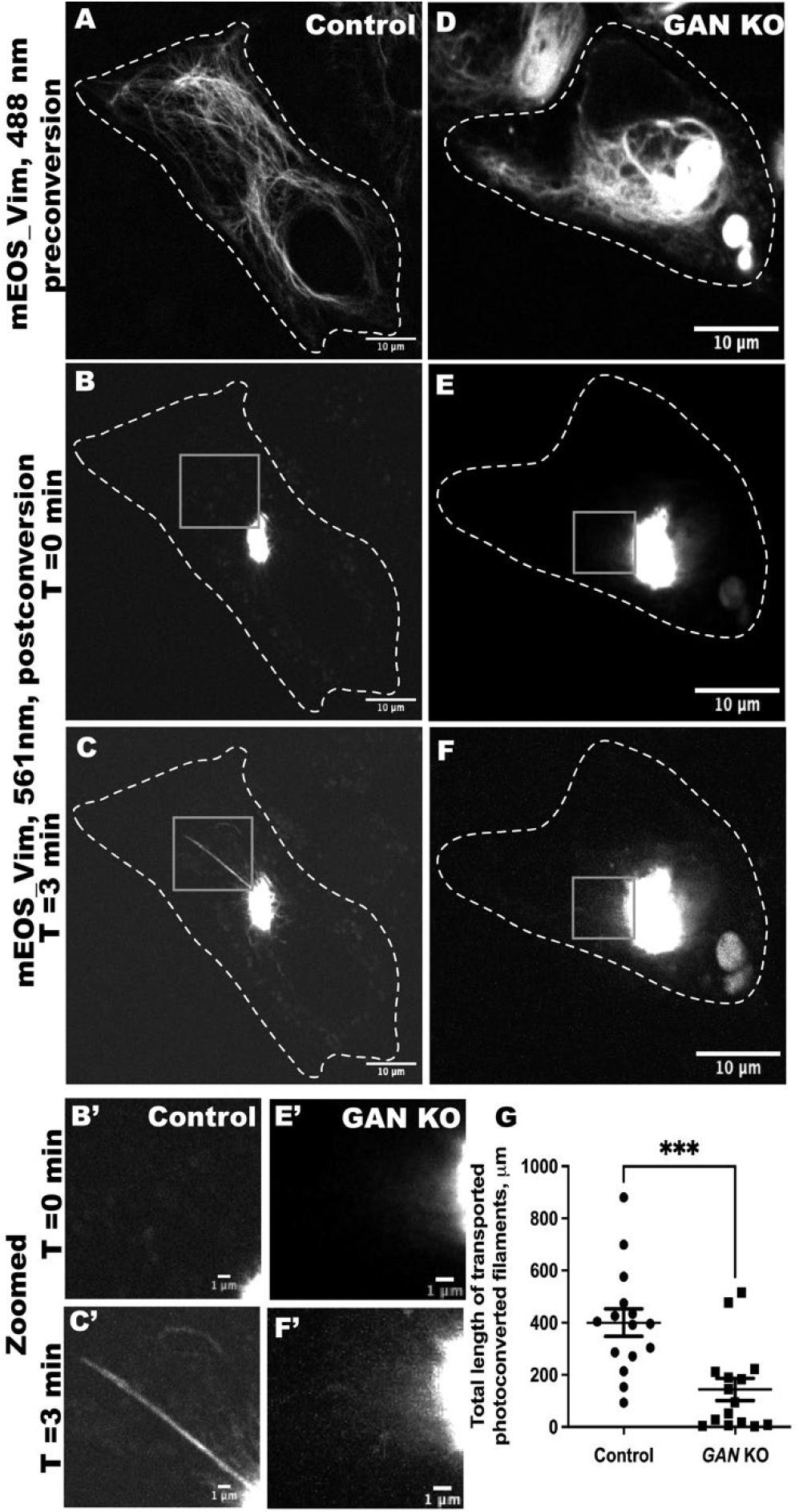
*GAN* KO inhibits the transport of vimentin filament. Photoconversion of mEOS-vimentin in RPE cells using spinning disk confocal microscopy in control (A-C) and *GAN* KO (D-F) cells. Panel A & D imaged under green channel (488nm) before photoconversion. Panel A shows vimentin filament in a control cell, while panel D displays the aggregated vimentin filament in a *GAN* KO cell. mEOS-vimentin was photoconverted from green to red at the specific region. Panels B & E were imaged under red channel (561nm) at time 0 min after photoconversion and C & F after 3 mins of photoconversion. Gray box region indicates the zoomed picture shown at bottom panel. Dotted lines mark the boundary of the cell. Scale bar 10 μm for full images and 1 μm for zoomed images. G) Photoconverted vimentin IFs outside the conversion zone were quantified and segmented filaments were counted for each frame. Statistical significance was determined using Student’s t-test (n = 15 cells). ***P, 0.005

To test whether this IF transport inhibition was specific for RPE cells, we used undifferentiated SH-SY5Y (human neuroblastoma) cells to test VIF transport in control and *GAN* knockdown conditions. *GAN* gene expression was knocked down using stable expression of an shRNA against the *GAN* gene (*GAN* KD) and cells expressing a scrambled shRNA served as a control. VIF transport was studied by transient expression of mEOS-vim. Control cells displayed well-developed and typically organized VIF meshwork (Sup Fig 1A). *GAN* KD cells displayed classic juxtanuclear aggregation of VIFs (Sup Fig 1D). VIF dynamics were measured by photoconversion of the EOS-vim as described above. We found that control cells displayed a robust transport of filaments from the photoconversion zone (Sup. Fig 1C, C’ and Video 5). In contrast, VIF transport in *GAN* KD cells was severely inhibited (Sup. Fig 1F, F’ and Video 6). Therefore, gigaxonin loss causes inhibition of IFs transport is not restricted to RPE cells only.

### Neurofilament transport is inhibited after *GAN* knock out

We are interested in studying NF dynamics after gigaxonin loss as NFs are primarily affected in GAN pathology. Neuroblastoma SH-SY5Y cells are used to study NF dynamics as they display a neuronal phenotype upon differentiation (27). mEOS_NFM was expressed in SH-SY5Y cells using transient transfection. Using retinoic acid in a low-serum medium, differentiation was induced 8 hours after transfection. Both control cells and *GAN* KD cells showed neurites 48 hours after differentiation was induced. The photoconversion approach was used to study NF transport. Control cells displayed proper NFM distribution (Fig 3A) in the cell body and neurites and active NF transport (Fig. 3C and C’ and video 3). Compared to control cells, *GAN* KD cells exhibit disorganized NFM and formed aggregates in the cell body (Fig 3D). Transport of NFs was significantly impacted in *GAN* KD cells, just like it was for VIFs (Fig. 3F and F’ and video 4). In *GAN* KD cells, quantification revealed significant inhibition of NFM transport (Fig 3G). Gigaxonin is not only required for VIF but also for NF.

**Figure 3:**
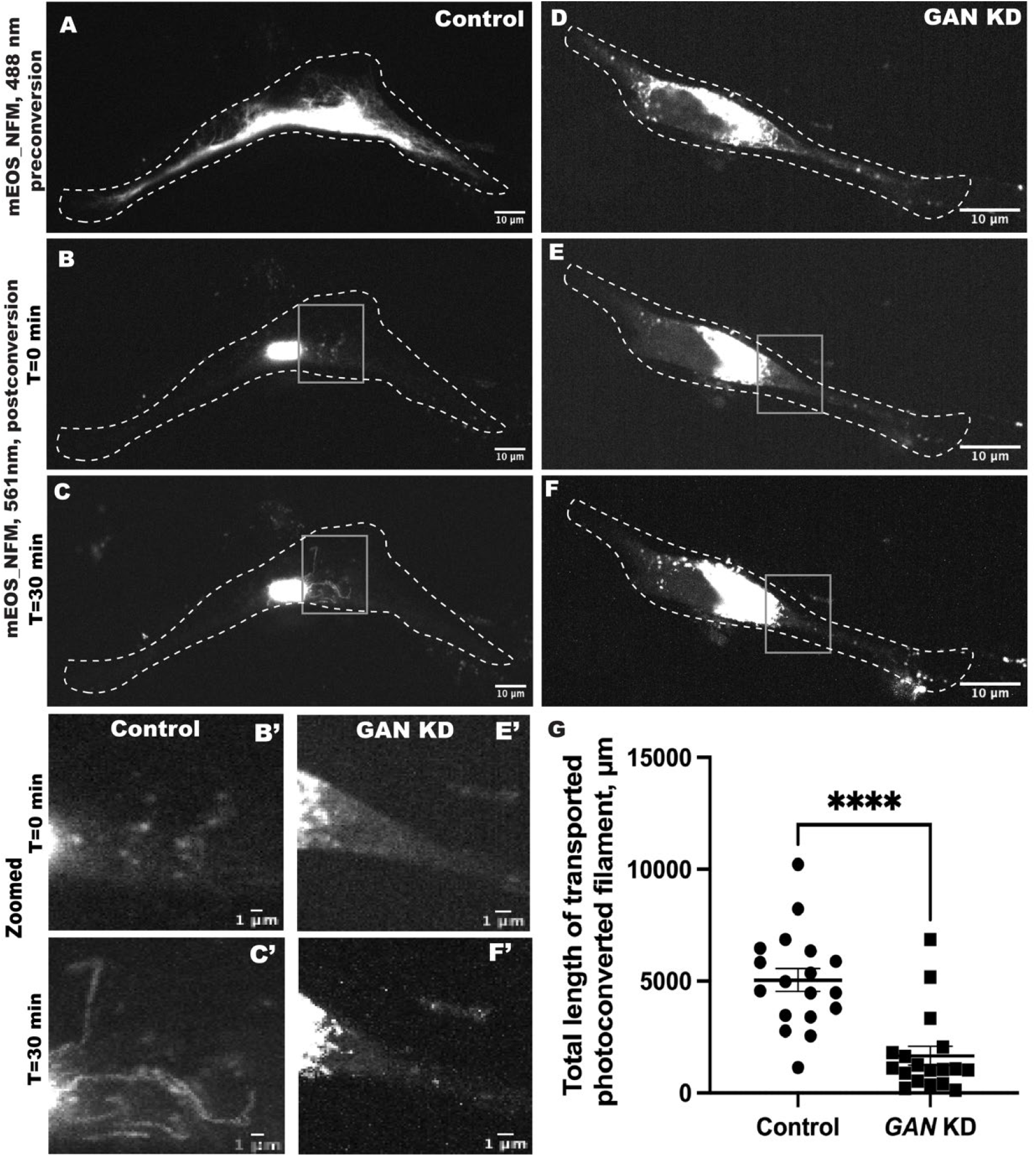
Neurofilament transport is inhibited in *GAN* KD cells. Photoconversion of mEOS-NFM in SH-SY5Y cells using spinning disk confocal microscopy in control (A-C) and *GAN* KD (D-F) cells. Panels A & D were imaged under the green channel (488nm) before photoconversion. Panel A shows the NFM filament in a control cell while panel D displays the aggregated NFM filament in *GAN* KD cells. mEOS-NFM was photoconverted from green to red at the specific region. Panels B & E were imaged under red channel at time 0 min after photoconversion and C & F after 30 mins of photoconversion. Dotted lines mark the boundary of the cell, gray box regions were zoomed and shown below. Scale bar 10 μm for full images and 1 μm for zoomed images. G) Photoconverted NFM IFs outside the conversion zone were quantified and segmented filaments were counted for each frame. Statistical significance was determined using Student’s t-test (n = 15 cells). ***P, 0.0005

### *GAN* KO increases the soluble fraction of vimentin

IF proteins exist as mature filaments and subunits/precursor oligomers, where the former is referred to as the insoluble, and the latter as the soluble fraction of cellular IFs (28, 29). In normal cells, the vast majority of vimentin is present in the insoluble filamentous form and only a small fraction is soluble (30). Earlier reports showed that the total IF protein level in *GAN* KO cells is increased without changes in the mRNA level (12, 31). Here we analyzed separately the soluble (IF precursors) and insoluble (mature IF) fractions of vimentin in RPE cells. Our results confirm that in *GAN* KO cells, the vimentin expression level was increased both in the soluble and insoluble fractions. While in the fraction of insoluble mature vimentin IFs increased modestly, in agreement with the previous reports, the amount of soluble vimentin oligomers was increased dramatically, about 20-fold (Fig 4).

**Figure 4:**
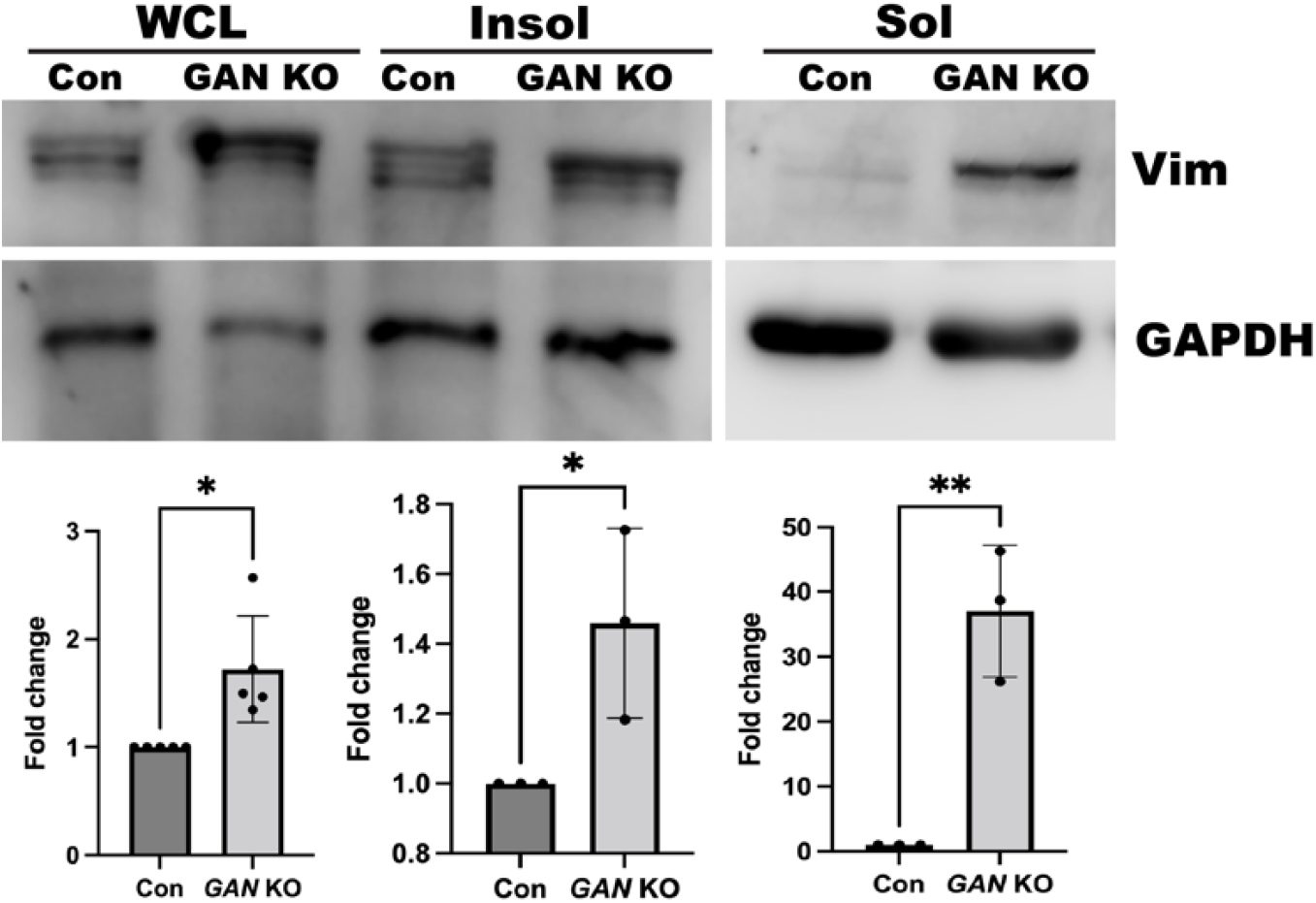
GAN KO cells show increased soluble vimentin. Immunoblotting analysis for vimentin in whole cell lysate (WCL), insoluble fraction (Insol) and soluble fraction (Sol). Vimentin protein level was increased both in WCL and insoluble of *GAN* KO cells. Soluble fractions show dramatic increase in vimentin in GAN KO cells. GAPDH protein levels were used as loading control. Fold changes are calculated with GAPDH normalized values.

### The kinesin-1 motor itself is functional in *GAN* KO cells

Lack of IF motility in *GAN* KO cells raised the question of whether gigaxonin is required for the function of kinesin-1 itself. We first tested if *GAN* KO affects expression of kinesin-1. Western blotting with a kinesin-1 heavy chain antibody demonstrated that the expression of the kinesin-1 protein was unaffected (Sup fig 2A). To test kinesin-1 functionally we studied the motility of lysosomes and autophagosomes in *GAN* KO cells to check for the ability of kinesin to transport cargoes unrelated to IFs. We found that lysosome (Fig 5A & B) and autophagosome (Fig 5 C & D) distributions in *GAN* KO cells were similar to their distributions in the control cells. Furthermore, after *GAN* KO there were no significant differences in the organelle motility between *GAN* KO cells and control cells. Normal movement of membrane organelles (lysosomes and autophagosomes) and normal distribution of microtubules (Sup fig 2E) in *GAN* KO cells, demonstrated that kinesin-1 motor was present and was fully functional and the cells contained many microtubule tracks.

**Figure 5:**
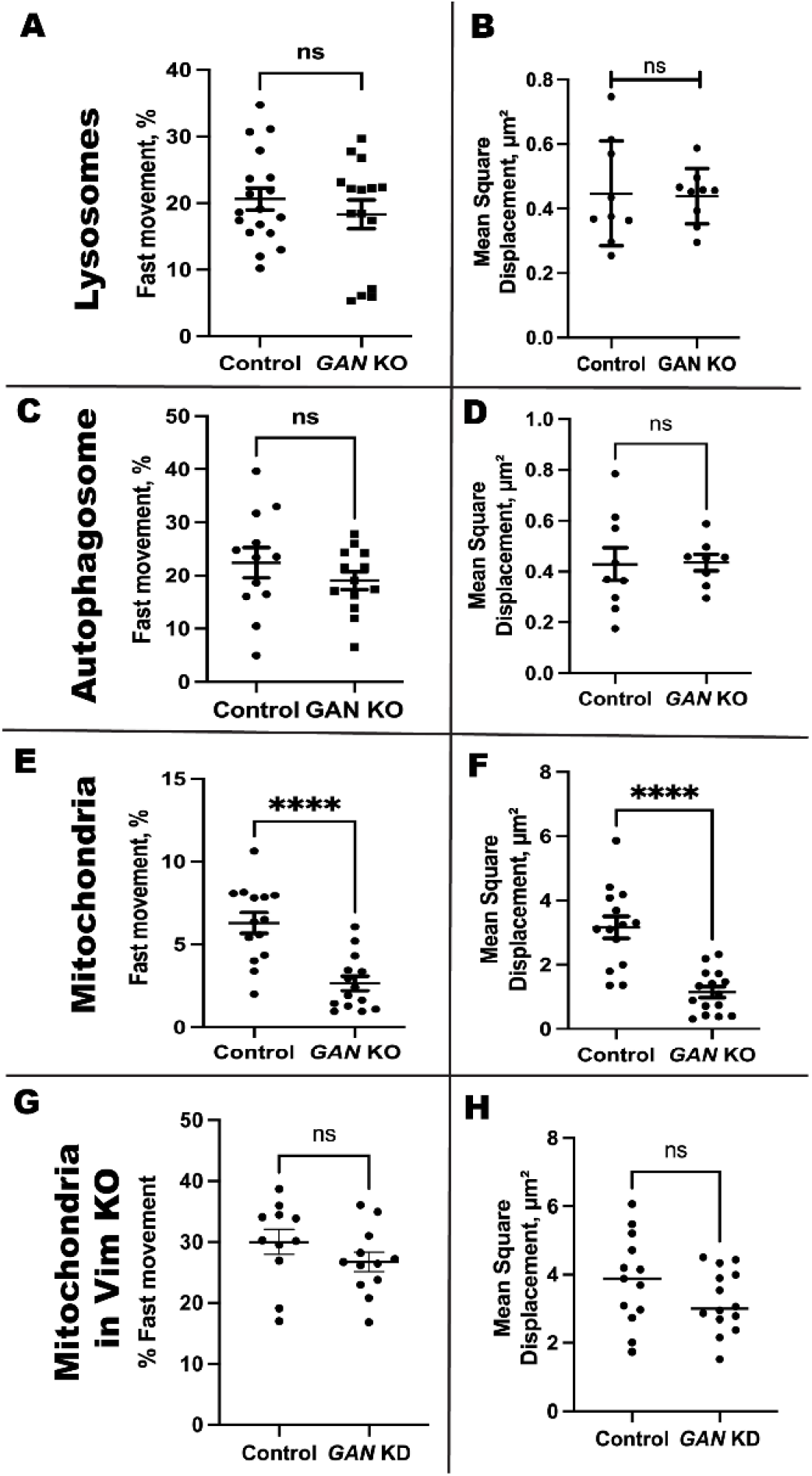
Membrane organelle motility under *GAN* KO and *GAN* KD. Lysosomes were visualized in live cell using lysotracker red. Panels A & B depict the fast movement and mean square displacement of lysosomes from control and *GAN* KO cells respectively. Autophagosomes were visualized in live cell using the autophagosome specific marker LC3 tagged with mCherry. Panels C & D the fast movement and mean square displacement of autophagosomes. Both lysosome and autophagosome motility are not affected in *GAN* KO cells. Mitochondria were visualized in live cell using Mitotracker red. (E) Fast mitochondrial movement in control and GAN KO cells, shown in percentage. *GAN* KO cells showed significant less mitochondrial motility. F) Mean square displacement of mitochondria in *GAN* KO is significantly lower that control cells. Silencing GAN expression in Vim KO cell has no effect on mitochondrial motility, fast motility (G) and mean square displacement (H).

### Vimentin accumulation traps mitochondria in GAN KO cells

Kinesin-1 is required for anterograde transport of mitochondrial (32, 33). Previous research has shown that the mitochondria in dorsal root ganglia neurons deficient of *GAN* gene are less motile and dysfunctional (31, 34). Here we studied the motility of mitochondria in *GAN* KO cells. In control cells, mitochondria were evenly distributed throughout the cell, while in *GAN* KO cells their distribution was severely altered. They were rarely seen at the cell periphery and mostly were localized close/within the VIF aggregate. In *GAN* KO cells, mitochondrial motility was significantly lower than in control cells. (Fig 5E). We also computed the mean square displacement, which revealed that *GAN* KO cells had significantly lower mean square displacement (Fig 5F). Loss of mitochondrial motility is consistent with the earlier studies. Although having functional kinesin-1 motors, mitochondrial motility was strongly inhibited in *GAN* KO cells.

As kinesin-1, the motor that moves mitochondria along microtubules is present and can move other membrane organelles, we hypothesize that the lack of mitochondria motility is explained by their binding to immotile IFs. To test this hypothesis, we combined KO of vimentin and silencing of *GAN*. Using *shRNA, GAN* gene expression was silenced in *Vim* KO cells, and the mitochondrial dynamics were investigated as previously mentioned. *GAN* silencing had no effect on the distribution or motility of the mitochondria in *Vim* KO cells (Fig 5 G & H). This result shows that the inhibition of mitochondria movement in *GAN* KO cells is a secondary effect of IF immobilization by VIF and the lack of VIF motility.

## Discussion

Giant axonal neuropathy (GAN) is a rare autosomal recessive, progressive neurodegenerative disorder with onset in early childhood and death usually by the second / third decade of life. GAN is caused by an abnormal or complete loss of function of gigaxonin (35). Aggregation of NFs has been reported in various neurodegenerative disorders like Parkinson’s syndrome, Alzheimer’s, and Amyotrophic Lateral Sclerosis (36, 37). However, GAN is a unique condition where all IFs undergo aggregation in all cell types including endothelial cells, epithelial cells, skin fibroblasts, muscle fibers, Schwann cells, and neurons (3, 26, 28). This indicates that gigaxonin plays an important role in IF regulation and organization. IFs form a complex network in the cytoplasm of mammalian cells. This network undergoes constant reorganization by movement, severing and, annealing to accommodate the constant need for changing cell shape and cellular environment (38). The organization of VIF and NFs is severely altered and filaments form juxtanuclear aggregates after the loss of the *GAN* gene as shown previously (25). Examination of platinum replicas of *GAN* KO cells revealed that individual filaments with normal diameter are present within this compact aggregate. This is in agreement with published data of GAN patient cells and cells with *GAN* KO in which individual NF and other IFs filaments with normal diameter can be observed (7, 12, 25, 34). The presence of individual filaments in GAN aggregates shows that they could be dynamic and transported. Hence, a failure of IF transport appears to be a possible explanation for their disorganization and aggregation in GAN.

In this work, IF transport was studied using a photoconversion technique. We observed that the dynamics of IFs (VIF and NF) were severely affected in the absence of gigaxonin protein. As IF anterograde transport is performed by the major microtubule motor kinesin-1 (23), we tested the possibility that inhibited transport of IFs could be explained by the general failure of kinesin-1. Strikingly, we found under *GAN* KO conditions, in addition to IF only mitochondrial motility was disrupted, whereas other organelles such as lysosomes and autophagosomes were unaffected. Taken together it is clear that kinesin-1 activity is not affected by gigaxonin loss. However, to address the mitochondrial dysfunction in GAN disorder, importance of IFs in mitochondrial anchoring and positioning should be considered (39). EM imaging showed mitochondria are associated with the IFs aggregate of *GAN* KO cells (12). In *GAN* KO cells, mitochondria exhibit altered metabolism and elevated oxidative stress (31). Loss of mitochondrial transport has been shown as the reason for distal axonal loss and atrophy in GAN (31). Poor neuronal development in a number of neurodegenerative disorders is associated with defective motility and function of mitochondria (40). In this study, we demonstrated how IF transport inhibition causes its meshwork collapse and aggregates in GAN. When IF becomes disorganized, mitochondrial activity and motility are impeded (34); moreover, IF traps the related mitochondria in its aggregate. So, in GAN, the inhibited IF transport may lead to mitochondrial dysfunction and play a role in pathogenesis. Since kinesin-1 is active in the *GAN* KO condition, there must be other reasons for the impeded transport of IFs.

Since gigaxonin is necessary for the regulation and efficient turnover of IFs, its absence or loss of function results in accumulation of IFs. Proteomic and biochemical approaches have shown that gigaxonin can bind to IFs directly (9, 12). Ectopic overexpression of gigaxonin results in the complete clearance of IFs from the cell over time (12). It has been proposed that mature IF filaments disassemble into soluble IF forms, which are gigaxonin substrates and are subsequently degraded (12, 41). Here we show dramatically increased levels of soluble vimentin with loss of gigaxonin. Thus, the primary outcome of gigaxonin loss is accumulation of soluble IF oligomers due to poor turnover. It is likely that IFs are connected to kinesin-1 by a yet unknown protein adapter. In GAN conditions, accumulating soluble IF oligomer may sequester this adaptor and thereby in a dominant-negative way inhibit IF transport (Fig 6). Consequently, the identification of the adapter that mediates the binding of kinesin-1 to IFs will allow us to directly test this hypothesis.

**Figure 6:**
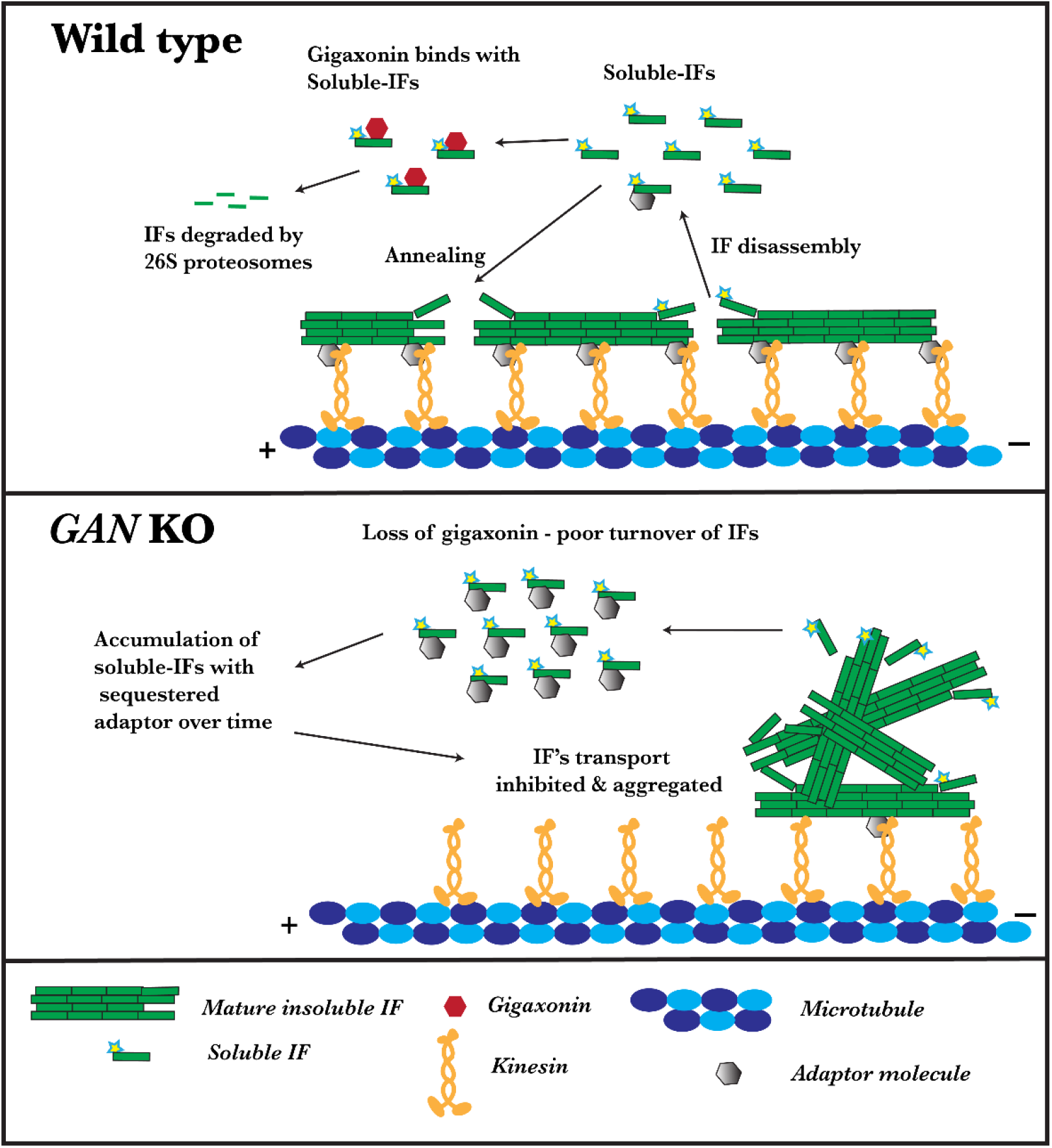
Loss of gigaxonin leads to inhibited transport of IF and its aggregation

## Materials and methods

### Plasmids

Vimentin tagged with mEOS3.2 photoconvertible tag was cloned in the pQCXIN vector as described (20). To generate pQCXIN mEOS3.2_NFM, vimentin in pQCXIN mEOS3.2_vim was replaced with NFM insert using BamHI and EcoRI digestion and ligation. Autophagosome-specific marker mCherry-hLC3B-pcDNA3.1 was a gift from David Rubinsztein (Addgene plasmid # 40827; http://n2t.net/addgene:40827; RRID: Addgene_40827)

### Cell culture, Transfection, and stable cell lines

All cell lines were maintained at 37 °C in 5% CO_2_. Human retinal pigment epithelial (RPE) cells were cultured in DMEM media (Sigma, #D5648) with 10% fetal bovine serum (FBS) (Neuromics, Minneapolis, MN, USA) supplemented with antibiotics (penicillin & streptomycin). RPE cells stably expressing mEOS3.2_vimentin (mEOS_vim) have been described previously (23). CRISPR was used to generate *GAN* KO in the RPE mEOS_vim cell line. Following the manufacturer’s protocol, the gigaxonin double nickase plasmid (Santa Cruz, #sc-407001-NIC) was stably expressed in RPE cells using retrovirus. The cells were selected with puromycin to obtain pure clonal populations of *GAN* KO cells. SH-SY5Y neuroblastoma cells were cultured in a 1:1 mixture of EMEM (BD, #670086) and F12 (Gibco, #11765-054), supplemented with 10% FBS and antibiotics (penicillin & streptomycin). Scrambled and *GAN* KO cell lines were generated by transducing SH-SY5Y cells with lentivirus expressing the Mission pLKO.1 backbone for scrambled shRNA (Sigma, #SHC002) and GAN shRNA (Sigma, TRCN0000083861) respectively. The transduced cells were selected with puromycin for pure clonal cell generation. For differentiation of SH-SY5Y cells into neurons, cells were incubated for 48h with media that contained 1% FBS and 100 μM all-trans retinoic acid (Thermo, #207341000).

For whole cell lysate preparation, cells were lysed using RIPA buffer [50mM Tris (pH7.4), 150mM NaCl, 1% TritonX-100, 0.5% sodium-deoxycholate, 0.1% SDS,] supplemented with peptidase inhibitor (Chymostatin, leupeptin, and pepstatin A, 20 μg/ml) and serine protease inhibitors (1 mM PMSF), then supernatant was collected after centrifugation at 10,000 x g for 10 min. For soluble and insoluble fraction preparation, we followed the protocol described earlier (29). Briefly, confluent 10 cm dishes were washed once with ice-cold PBS and the cells were lysed in a buffer containing 0.5% Triton X-100, 150 mM NaCl, 20 mM Tris·HCl pH 7.4, 2 mM EGTA, 2 mM EDTA, 1.5 mM sodium vanadate. Then cell lysates were ultracentrifuged for 30 min at 265,000 × g and the pellet (insoluble fraction) and supernatant (soluble fraction) were collected. Acetone precipitation was used to concentrate the protein that is soluble. In brief, two volumes of ice-cold acetone were added to the soluble fraction and incubated at -20 °C for 2 h. After centrifugation at 10000 x g for 10 min, the pellet was reconstituted in 5X Laemmli buffer and Western blot analysis was carried out. For the Western blot analysis, equal protein loads were maintained among samples. Antibodies used in these studies included chicken polyclonal anti-vimentin (Biolegend, Cat #919101, dilution 1:2000) and HRP-conjugated mouse monoclonal GAPDH (Proteintech, cat HRP-60004, dilution 1:10000). Blots were developed using WesternBright ™ Quantum (Cat # K-12042-D20) and imaged using Li-COR Odyssey Fc imaging instrument with Image studio software version 5.2.

### Live cell imaging

Tokai-Hit stage-top incubator (Tokai-Hit) and Okolab gas mixer (Okolab) were used to maintain temperature at 37 °C and CO_2_ at 5% throughout the live imaging. Images were collected using a Nikon Eclipse Ti2 stage equipped with a W1 spinning disk confocal head (Yokogawa CSU with pinhole size 50 μm), 60X 1.49 N.A. or a 100X 1.45 N.A. oil immersion lenses. Photometrics Prime 95B sCMOS or Hamamatsu 29 ORCA-Fusion Digital CMOS Camera driven by Nikon Elements software were used for image acquisition.

We have demonstrated previously that cytoskeletal dynamics can be studied efficiently by fluorescence microscopy of live cells using photoconversion of mEOS fluorescent protein fused with a cytoskeletal protein of interest (20). Photoconversion of mEOS3.2 was achieved using 405 nm illumination from a Heliophor 89 North LED mounted in the epifluorescence pathway. A pinhole diaphragm was included in the light path for confinement of the photoconversion zone. For vimentin dynamics study, cells were plated on glass coverslips one day before imaging. Cells expressing mEOS_Vim were photoconverted for 3 s and then timelapse sequences were acquired for 3 min with the interval of 15 s, using 561 nm laser excitation. Neurofilament dynamics were studied in differentiated SH-SY5Y cells. Undifferentiated SH-SY5Y cells were electroporated with mEOS_NFM in pQCXIN plasmid. The electroporated cells were plated in complete medium onto coverslips coated with poly-L-lysine. Neuronal differentiation was induced 8h after electroporation, by incubating cells with 10 μM all-trans retinoic acid in the low serum medium (1% of FBS). After 48h of differentiation, NFM dynamics was recorded for 30 min with interval of 1 min after photoconversion. For mitochondrial and lysosomal motility 25 nM MitoTracker Red CMXRos (Invitrogen, #M7512), and 25 nm lysotracker red (Invitrogen, #L7528) were used respectively. Live cells where incubated for 10 mins with the respective dyes and then imaged. For autophagosome analysis, cells were transiently transfected with the autophagosome specific marker protein LC3B tagged with mCherry. Autophagosomes were induced 24h post-transfection using Earls Balanced Salt Solution (EBSS) buffer, pH 7.4 for 3h. Organelles’ motilities were captured for the total time of 2 – 3mins with interval of 2 – 5 seconds.

### Image analysis

Fiji ImageJ plugins were used to analyze IF transport (42). To determine the overall movement of photoconverted filaments, we used the curve tracing plugin (43). Photoconverted filaments which transported outside the initial photoconverted region were only taken for quantification and the photoconversion zone was masked in all frames while quantifying the transported filaments. To get the overall length of filaments transported over the duration of imaging, we calculated the total length of filaments transported for each frame and added all the values. Our plugins are deposited in GitHub, in the following URL (https://github.com/mkitti/CurveTracingUtils).

The analysis explorer package in Nikon Elements software (NIS elements AR, version 5.30.03) was used to compute the motility of mitochondria, lysosomes and autophagosomes. Preprocessing includes background correction and smoothing after which the particles were analyzed by the random motion method. Organelles that appeared in more than five frames were taken into consideration for analyzing the motility, and non-motile organelles were excluded from the analysis. We had a velocity threshold of 200 nm/s for fast-moving organelles.

### Immunostaining

Fixation and staining protocols were performed as described previously (23). Cells were incubated with primary antibodies for 1h at room temperature. Primary antibody used in this study was microtubule (Dm1α mouse monoclonal). Secondary antibodies used in this study were Rhodamine conjugated goat anti-mouse (Jackson ImmunoResearch, #115-025-003), Alexa-Fluor 647-conjugated donkey Anti-Mouse (Jackson ImmunoResearch, #715-605-151), and Rhodamine (TRITC) goat anti-rabbit antibody (Jackson ImmunoResearch, #111-025-003). For actin staining, cells were stained with rhodamine-conjugated phalloidin (Thermo Fisher Scientific, #R415) for 1h before mounting.

### Electron microscopy of platinum replicas

Cells were grown overnight on glass coverslips. The membrane and soluble cytosolic proteins were extracted with 1 % Triton-X 100 in PHEM buffer (80mM Pipes, 1mM EGTA, 1mM MgCl_2_ at pH 6.8) for 5 min. Cells were treated with 0.6 M potassium chloride in PHEM buffer, for 10 min to remove microtubule and actin filaments from the cytoskeletons still attached to the coverslips. Platinum replica electron microscopy (PREM) was performed as previously described (44, 45). Briefly, samples were fixed with 2% glutaraldehyde in 0.1M cacodylate buffer, tannic acid, and uranyl acetate; critical point dried; coated with platinum and carbon; and transferred onto electron microscopic grids for observation. Samples were imaged using a FEI Tecnai Spirit G2 transmission electron microscope (FEI Company, Hillsboro, OR) operated at 80 kV. Images were captured by Eagle 4k HR 200kV CCD camera.

### Statistical analysis

Data were analyzed using GraphPad Prism 9.3 (GraphPad Software Inc.) and are shown as mean with SEM. Statistical testing involved unpaired Student’s t tests. Error bars indicate 95% CI.

## Abbreviation

IF: Intermediate filament
VIF: vimentin intermediate filament
NF: Neurofilament
GAN: Giant axonal neuropathy
KO: Knock out:

## Acknowledgement

Authors thank Dr. Stephen Adam for stimulating discussions and critical reading of the manuscript and Dr. Farida Korobova, Center for Advanced Microscopy (Northwestern University) for PREM sample preparation and EM image acquisition and This work is supported by the National Institute of General Medical Sciences (NIH) under grants P01 GM096971 and R35 GM131752. Center for Advanced Microscopy is supported by NCI CCSG P30 CA060553 awarded to the Robert H Lurie Comprehensive Cancer Center.

