## Supplementary information for "Gigaxonin is required for intermediate filament transport"

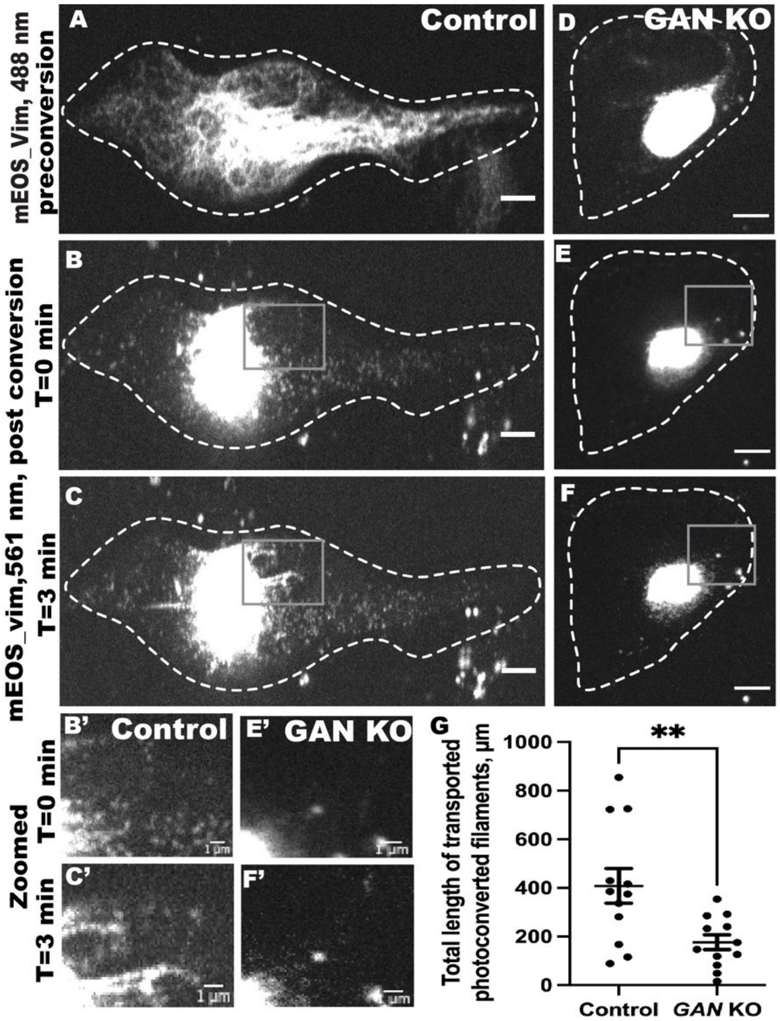


**Supplementary figure 1:** Vimentin IF transport is inhibited in undifferentiated SH-SY5Y *GAN* silenced cells. Photoconversion of mEOS_vim in undifferentiated SH-SY5Y cells using spinning disk confocal microscopy in control (A-C) and *GAN* KO (D-F) cells. Panel A & D imaged under green channel (525nm) before photoconversion. Panel A shows vimentin filament in a control cell, while panel D displays the aggregated vimentin filament in a *GAN* KO cell. mEOS-vimentin was photoconverted from green to red at the specific region. Panels B & E were imaged under red channel at time 0 min after photoconversion and C & F after 3 mins of photoconversion. Gray box region indicates the zoomed picture shown at bottom panel. Dotted lines mark the boundary of the cell. Scale bar 5 μm for full images and 1 μm for zoomed images. G) Photoconverted vimentin IFs outside the conversion zone were quantified and segmented filaments were counted for each frame. Statistical significance was determined using Student's t-test (n = 15 cells). ***P, 0.0005

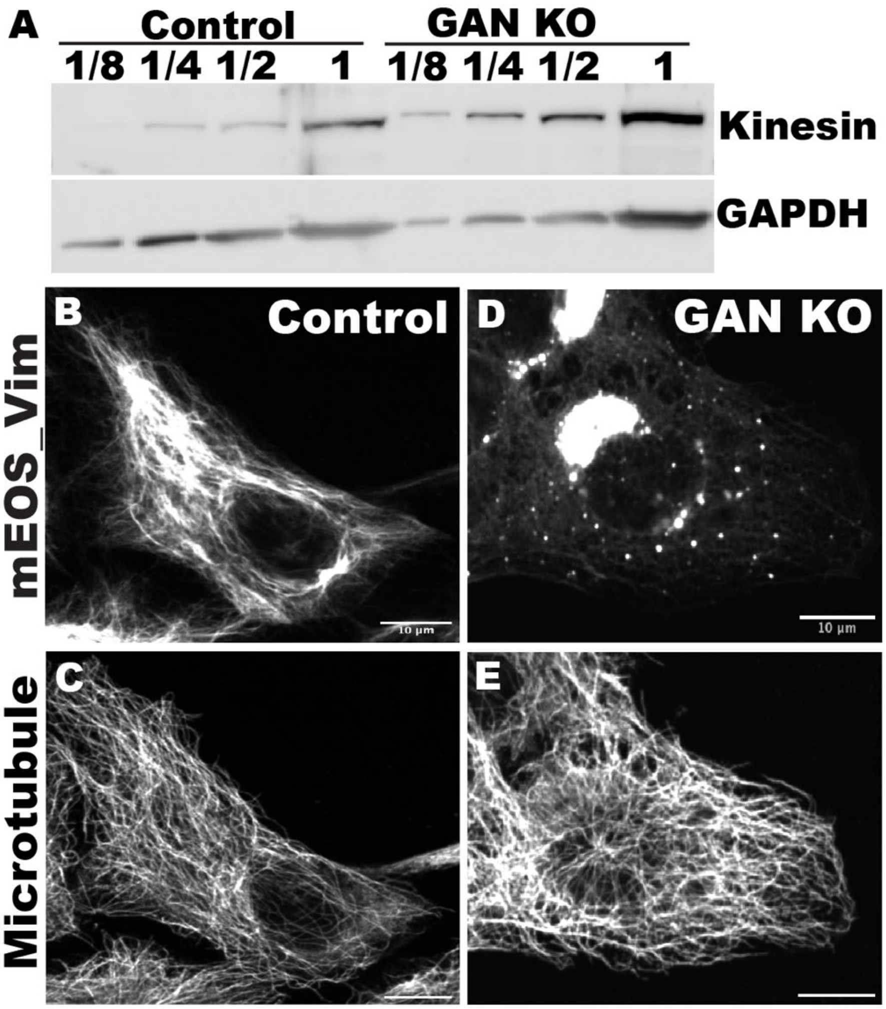


**Supplementary figure 2**: Kinesin expression and microtubule organization were not affected in *GAN* KO cells. A) Western blot showing the kinesin expression in whole cell lysates of control and GAN KO cells. Immunostaining of microtubule organization, right panel shows the normal distribution of vimentin (B) and microtubule (C) in control cell. Left panel shows aggregated vimentin (D) and normally distributed microtubule (E) in *GAN* KO cells

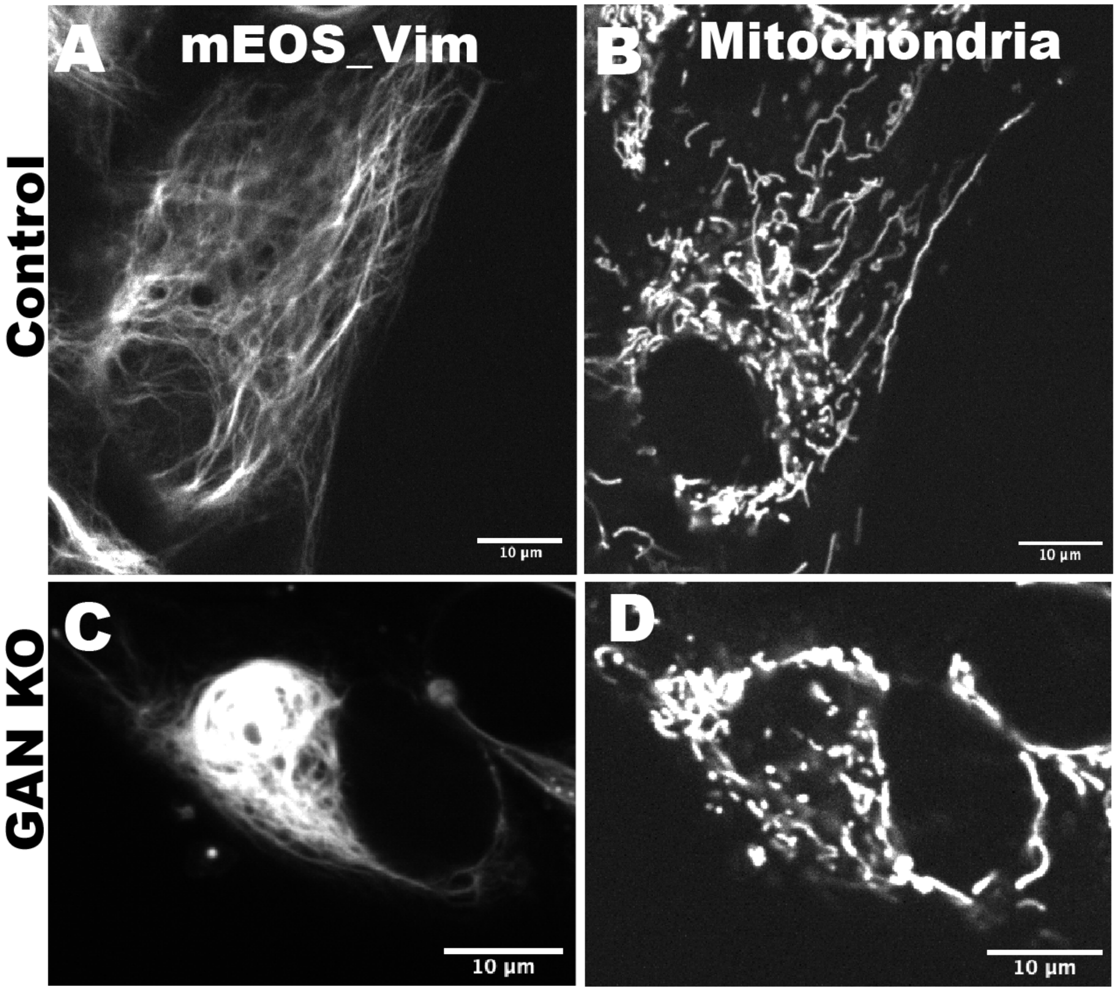


**Supplementary figure 3:** Distribution of mitochondria is affected in *GAN* KO cells. (A) Shows the vimentin distribution in control cells and (C) shows the aggregated vimentin in *GAN* KO cells. Mitochondria is visualized using MitoTracker dye. B) Mitochondria in control cells is distributed all over the cell and in (D) *GAN* KO cells mitochondria is concentrated and localized with vimentin aggregate.

**VIDEO CAPTIONS**

**Video 1.** Vimentin filaments are transported in RPE mEOS_Vim cells. A 10 µm-diameter area of a mEOS_Vim was photoconverted from green (488 nm) to red (561 nm) with 405nm light (see gray circle). The movie shows the transport of red vimentin filaments outside of the photoconverted area every 15 sec after photoconversion for 3 min.

**Video 2.** *GAN* KO in RPE mEOS_vim inhibits the vimentin filament transport. A 10 µm-diameter area of a mEOS_Vim was photoconverted from green (488 nm) to red (561 nm) with 405nm light (see gray circle). The movie shows vimentin filaments are confined inside the photoconverted area, imaged every 15 sec after photoconversion for 3 min.

**Video 3.** *GAN* KO in RPE mEOS_vim inhibits the single vimentin filament transport. A 10 µm-diameter area of a mEOS_Vim was photoconverted from (488 nm) to red (561 nm) with 405nm light (see gray circle). Here we photoconverted the region of cell where single filament are present. The movie shows vimentin filaments are confined inside the photoconverted area, imaged every 15 sec after photoconversion for 3 min.

**Video 4.** NFM filaments are transported in differentiated SH-SY5Y cells. A 10 µm-diameter area of a mEOS_NFM expressing cell was photoconverted from (488 nm) to red (561 nm) with 405nm light (see gray circle). The movie shows the transport of red NFM filaments outside of the photoconverted area every 1 min after photoconversion for 30 min.

**Video 5.** GAN KD inhibits the NFM transport differentiated SH-SY5Y cells. A 10 µm-diameter area of a mEOS_Vim was photoconverted from (488 nm) to red (561 nm) with 405nm light (see gray circle). The movie shows NFM are confined inside the photoconverted area, imaged every 1 min after photoconversion for 30 min.

**Video 6.** Vimentin filaments are transported in undifferentiated SH-SY5Y. ells. A 10 µm-diameter area of a mEOS_Vim was photoconverted from (488 nm) to red (561 nm) with 405nm light (see gray circle). The movie shows the transport of red vimentin filaments outside of the photoconverted area every 15 sec after photoconversion for 3 min.

**Video 7.** GAN KD inhibits the vimentin filament transport undifferentiated SH-SY5Y. A 10 µm-diameter area of a mEOS_Vim was photoconverted from (488 nm) to red (561 nm) with 405nm light (see gray circle). The movie shows vimentin filaments are confined inside the photoconverted area, imaged every 15 sec after photoconversion for 3 min.
